# Serotype-based evaluation of an optogenetic construct in rat cortical astrocytes

**DOI:** 10.1101/2021.09.17.459681

**Authors:** Lakshmini Balachandar, Diana Borrego, Jorge Riera Diaz

## Abstract

Optogenetics is a modern technique which has been recently expanded to non-neuronal cell types, e.g., astrocytes, and involves targeted gene delivery of light-sensitive ion channels like Channelrhodopsin-2 (ChR2). Optogenetic regulation of astrocytic activity can be used for therapeutic intervention of several neurological disorders. Astrocytic gene delivery, viz adeno-associated viral (AAV) vectors, have proven to be robust, time-, and cost-efficient contrary to the generation of transgenic animal models. When transducing astrocytes with an AAV vector, it is imperative to perform a serotype evaluation of the AAV vector due to variability in serotype transduction efficiency depending on species, target region and construct length. Rats have been a very successful animal model for studying a variety of brain disorders, from which ChR2-based intervention of astrocytes will benefit. However, the most efficient AAV capsid serotype targeting astrocytes for ChR2 expression in the *in vivo* rat brain cortex has not been characterized. To address this, we have evaluated AAV serotypes 1, 5, and 8 of the vector AAV-GFAP-hChR2(H134)-mCherry targeting astrocytes in the rat brain neocortex. Results show that serotype 8 exhibits promising transduction patterns, as it has demonstrated the highest tangential and radial viral spread in the rat brain. Our research will facilitate translational research for future applications of optogenetics involving the transduction of rat brain cortical astrocytes.

## 1. Introduction

Astrocytes—a pivotal cell type in the brain, play key roles in neurovascular coupling [1] and regulation of neuroinflammatory responses [2]. Dysregulation of astrocytic function is observed in various disorders like Alzheimer’s disease, autism spectrum disorders, Parkinson’s disease and epilepsy [3]. Control of astrocytic dynamics can be critical in the intervention of these disorders. Optogenetics—a novel state-of-the-art technique provides a platform for precise cellular targeting and control using light, with the potential for delving into a variety of pathological conditions in the brain [4, 5]. One of the earliest investigations for astrocytic control using optogenetic methodologies by Gradinaru *et al* [6] demonstrated the relevance of thalamic stimulation *in vivo* by employing a lentivirus-based delivery of the optogenetic construct for probing astrocytes. Few studies in literature also point out to the investigation of astrocytic Ca^2+^ dynamics elicited by light stimulation, *in vitro* [7, 8] and recent transgenic approaches to study the role of astrocytic contribution to cerebral blood flow and neurovascular coupling, *in vivo* [9].

Delivery of the optogenetic construct targeting a specific cell type (i.e., astrocytes) can be achieved via two major approaches—transgenic animal model creation and through a viral vector. Methodologies involving the generation and maintenance of transgenic lines are time-consuming and costly [9, 10] while viral delivery of the construct has precise cellular targeting, is both time and cost-effective, and facilitates application of optogenetics to existing rat disease models [11]. For a multitude of reasons, adeno-associated viral (AAV) vectors are preferred for gene delivery over other viral vector backbone types—lentiviral (LV), adeno-viral vectors (AVV), and retroviral (RV). Their characteristics of low immunogenicity, ability to efficiently target both dividing and non-dividing cells, robust gene delivery and persistence of expression in rodent models make them potent tools for research and gene therapy. Furthermore, they can be engineered to facilitate high transduction capabilities of a diverse set of target tissues and species [12–14].

Cellular targeting using AAV-based optogenetics requires an appropriate selection of the serotype. AAV capsid serotype plays a crucial role in determining the efficacy and extent of gene delivery and transduction. Serotype-based delivery differs based on target cell type, brain area, species of intervention, and size of the construct which imparts unique capabilities of targeting a various range of cells. Tailoring the serotype also influences AAV vector transport and optimizing vector-host interaction [15–17]. Therefore, it is imperative to perform a comprehensive evaluation of an optogenetic construct for targeting a given brain region, cell type and species.

This study seeks to provide a holistic characterization of AAV serotypes of a ChR2-based optogenetic construct for targeting rat cortical astrocytes and achieving maximal stable transduction patterns. We evaluate transduction patterns of the optogenetic component—AAV-GFAP-hChR2(H134R)-mCherry in rat cortical astrocytes. The selection of potential serotypes (1, 5 and 8) used in this study were based on previous findings *in vivo* and *in vitro* for non-optogenetic constructs, in various CNS regions and species [18–20]. The virus was delivered to the rat brain by intracerebral injections and following perfusion and histology, astrocytic expression of the construct was evaluated. Upon quantification of transduction, we found that serotype 8 of the optogenetic construct had the maximum viral spread (in both tangential and radial directions), as compared to serotypes 1 and 5, in the astrocytes of the rat neocortex. Our results therefore suggest that rAAV8 can be employed as a tool for manipulating gene expression in rat cortical astrocytes *in vivo* for ChR2-based optogenetic interrogation. Our study provides an insight into a systematic serotype evaluation of AAV-based optogenetic astrocytic targeting which can further be used to establish robust astrocytic control and mediation for therapeutic interventions in neurological disorders.

## 2. Materials and Methods

### 2.1. AAV viral vectors

AAV vectors (AAV-GFAP-hChR2(H134R)-mCherry) of serotypes 1,5 and 8 were provided by The University of North Carolina Vector Core at Chapel Hill (~10^12^ GC/ml).

### 2.2. Animal preparation and stereotaxic injections

Male Wistar rats (7-10 weeks) were housed in standard cages at a 12h-12h light-dark cycle with free access to food and water. All experimental procedures and animal care in this study are approved and carried out in compliance with the Institutional Animal Care and Use Committee (IACUC) at Florida International University (Approval No. 16-032), IBC (16-001) and NIH guidelines. The rats were anesthetized using isoflurane anesthesia (5% for initial induction, 1.5% for maintenance, 1L/min O_2_, 14.7 PSI). After mounting the rat’s head on the stereotaxic frame (Narishige, Japan), the body temperature was maintained at ~37 °C using a water-circulating heating pad (TPZ-0510EA, Texas Scientific Instruments, LLC) with a pump (TP700, Texas Scientific Instruments, LLC). The temperature and respiration rate were monitored continuously using the PowerLabs 8/35 data acquisition device and the LabChart software (AD Instruments). An incision was made in the middle of the scalp and connective tissue was cleared to better visualize the *lambda* and *bregma* stereotaxic coordinates. The measurements were taken from the top of the skull, bregma and the midline. A burr hole was made at the injection coordinates—2 mm distal to the bregma and 2 mm from the midline, in the right hemisphere. 1 μL of the virus stock was injected at 50nl/minute, 1 μm into the cortex, using a microinjection syringe: Hamilton (80314: 10 μL, Model 701 RN SYR, 32 ga, 2 in, point style 3 needle), controlled by an UltraMicroPump with SYS-Micro4Controller (World Precision Instruments) and MPC-325 Multi-Micromanipulator system (Sutter Instruments). To prevent backflow of the viral volume, the needle was kept in place for 10 minutes after injection, withdrawn halfway and allowed to rest for another 10 minutes before being full withdrawn [21]. The scalp was sutured using silk braided surgical sutures (Keebomed). The rats were treated with Enroflox 100 (Norbrook) and Carprofen (Carprieve, 50mg/ml) for a week post-surgery to promote recovery, as well as to prevent infection and inflammation.

### 2.3. Tissue preparation

Two weeks after surgery, the rats were deeply anesthetized under isoflurane anesthesia (5%, 1 L/min O2, 14.7 PSI) and transcardially perfused with 4% paraformaldehyde. Subsequently, the brain was removed and placed in 4% PFA for 24 hours at 4°C. It was then transferred to 30% sucrose/PBS for cryoprotection at 4°C. The target region of the brain was embedded in Leica freezing medium and frozen at −80 °C. Coronal brain sections (30 μm thick) were made using a Leica cryostat (CM1950), mounted on Fisher Superfrost Plus slides and left to dry overnight, before cover slipping with Diamond Antifade Mountant (ThermoFisher Scientific) and imaging (refer to Fig. 1 for the stepwise procedure).

**Fig. 1.**
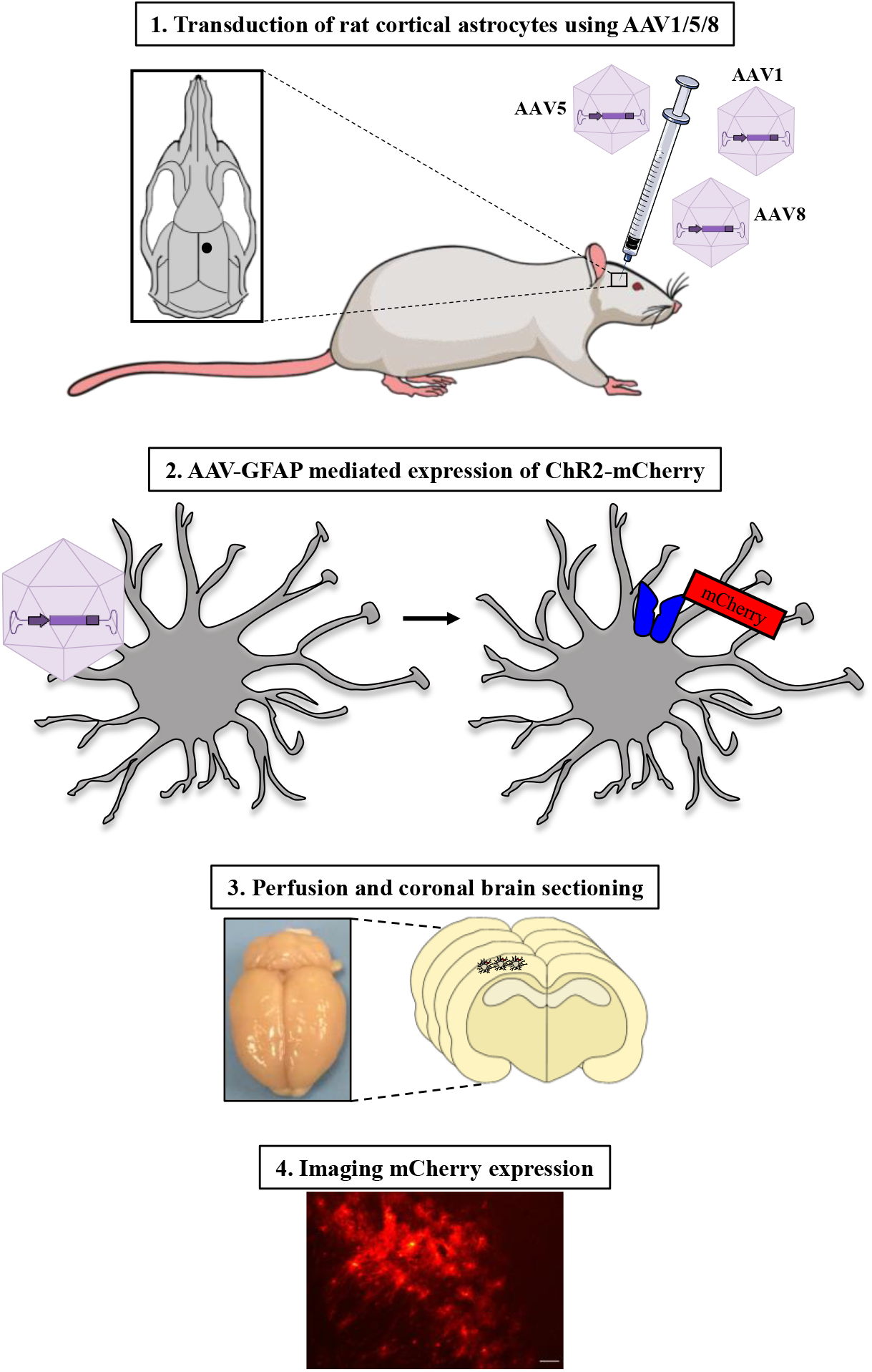
Experimental flow for studying rat cortical astrocyte transduction using various viral vector serotypes. Experimental steps **i**ncluding: 1. Microinjection of serotypes 1,5 or 8 of the AAV-GFAP-ChR2(H134R)-mCherry virus targeting rat cortical astrocytes. Inset shows microinjection location (2mm posterior to the Bregma, 2mm from the midline). 2. Targeted expression of ChR2-mCHerry on the GFAP^+^ astrocyte. 3. Perfusion and coronal brain sectioning, 2 weeks post-surgery and 4. mCherry expression analysis to determine serotype expression efficacy.

### 2.4. Quantitative analysis

To determine the viral spread of the various serotypes of the optogenetic construct AAV-GFAP-hChR2(H134R)-mCherry targeting astrocytes, we performed a quantification after intracranial injections of the virus (Sample size (n) = 5 rats/ serotype). The number of slices displaying mCherry expression were multiplied by a factor of 30 (indicative of the slice thickness of 30 μm) to obtain the final distance of spread of the virus in the brain. Transduced area per section was measured using FIJI [22]. Statistical tests were performed using GraphPad Prism version 8.4.3. The summary data is presented as mean ± std.

## 3. Results

### 3.1 Viral spread of serotypes of AAV-GFAP-hChR2(H134R)-mCherry in the adult rat brain neocortex

Our experiments showed that there was a significant difference between serotypes 8, 5 and 1 (Fig. 2), where serotype 5 had the least viral spread as compared to the others. After confirming normality of the data using the Shapiro-Wilkins test, a one-way ANOVA and Tukey’s multiple comparisons test was conducted to establish statistical significance. To determine the radial spread of the serotypes of the optogenetic construct, we employed ImageJ to calculate the area of transduction. Traces of the area of the central slice of expression were drawn and quantified on the software. From our experiments, we observe that serotype 8, had the highest radial spread of the virus in the rat brain cortex, as compared to both serotype 1 and 5 (Fig. 3). Thus, for targeting astrocytes optogenetically using this construct, rAAV8 had the maximum transduction efficiency. The results were obtained by performing a Shapiro Wilkins test for normality, non-parametric ANOVA with a Kruskal Wallis test, followed by a Welch’s T test between pairs of serotypes (significance declared at p<0.05).

**Fig. 2.**
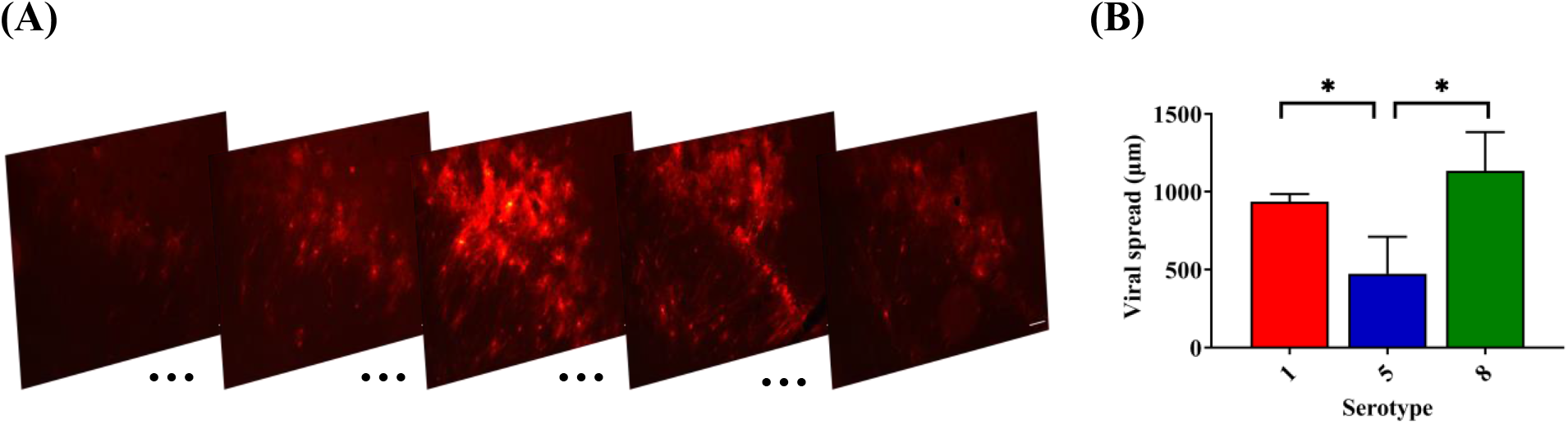
Quantification of serotype-based mCherry expression in the rat neocortex across coronal sections. **(A)** Representative images of AAV-GFAP-ChR2(H134R)-mCherry serotype 8 expression at 1) 1.69mm, 2) 1.78mm, 3) 2.02mm, 4) 2.29mm and 5) 2.41mm from the Bregma. **(B)** Quantification of the viral spread of the serotypes 1,5 and 8 (n = 5/ serotype). Average data are represented as mean ± std. 1-way ANOVA, p<0.001, 95% CI. Tukey’s multiple comparisons test reveals the significance of pairwise comparisons to be: Serotype 1 vs. 5 (**), Serotype 8 vs. 5 (***), Serotype 1 vs 8 (ns). Scale bar – 100μm.

**Fig. 3.**
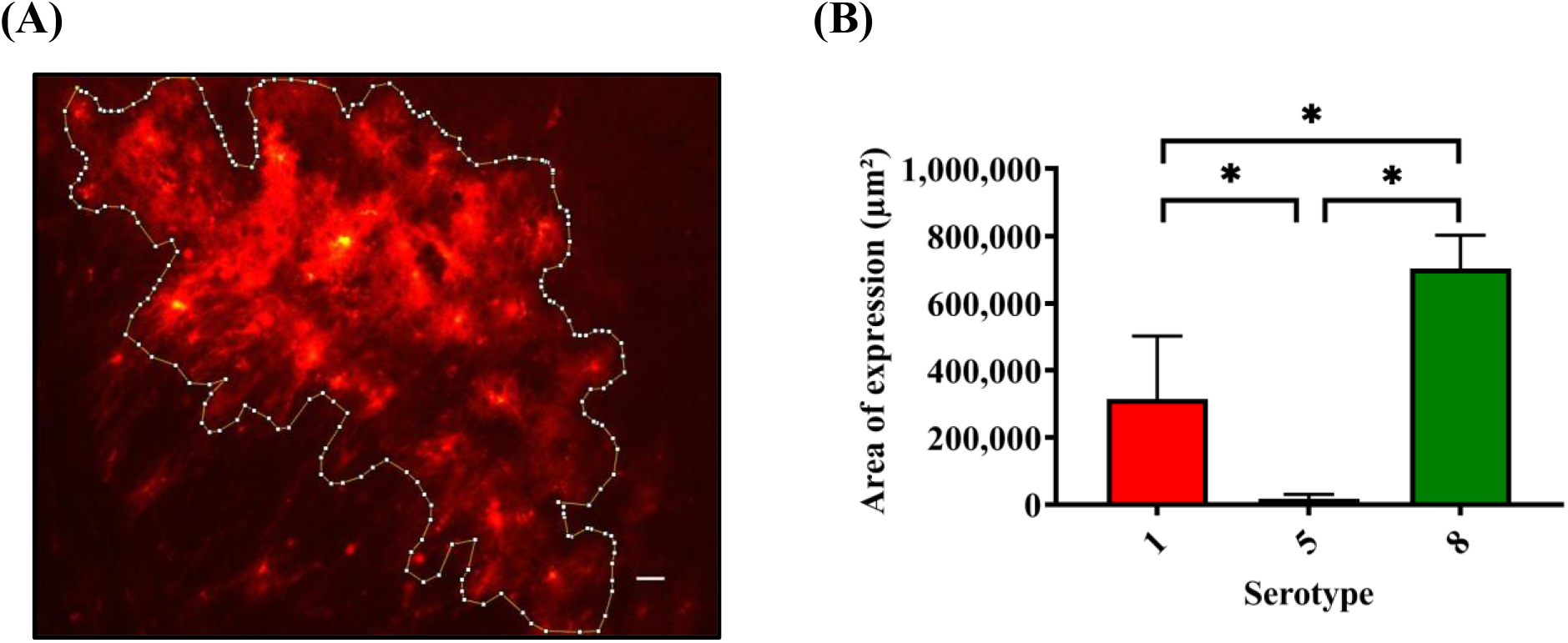
Quantification of radial expression of various AAV serotypes in the rat neocortex. **(A)** Representative image of an AAV8-transduced rat slice (enclosed within the highlighted bound area) **(B)** Quantification of the radial viral spread of the serotypes 1,5 and 8 (n = 5/ serotype). Average data are represented as mean ± std. Kruskal Wallis test, p<0.0001, 95% CI. Mann Whitney test reveals the significance of pairwise comparisons to be: Serotype 1 vs. 5 (*), Serotype 8 vs. 5 (**), Serotype 1 vs 8 (*). Scale bar – 100μm.

## 4. Discussion

AAVs are single-stranded non-enveloped DNA viruses with capsids, sized ~ 4.7 kb, making them among the smallest known viruses. They have been studied extensively to probe into their capabilities of gene delivery and therapy for neurological disorders [23, 24] with transduction efficacy varying greatly with each capsid, as detailed in their serotype. Studies suggest that transduction efficiency depends on the compatible serotype, target transduction area, and route of administration [25, 26]. The evaluation of tropisms of AAV capsid serotypes has been performed in a plethora of organs in a variety of rodent models. Specifically, serotypes 2, 4 and 5 have transduced ependymal cells [27], while, 7, 8, 9 and rh10 transduced neurons [25] via intracerebral injections in mice efficiently. Furthermore, serotypes 1, 2, 5 transduced the SN pars compacta showing high efficiency, additionally 1 and 5 transduced the pars reticulata in the CNS of rats [28]. Notably, even if the vectors contained the same genome, they exhibited markedly different patterns of transduction, owing to their serotype [16]. A study by Yang *et al* [29] details the difference in uptake of the genetic material via AAV delivery to the murine retina and elucidates how a shorter construct of the AAV2 serotype, has a significantly higher uptake by the retina, as compared to a longer construct of AAV2.

Our study indicates that for optogenetic targeting of cortical astrocytes, serotype 8 had the most efficient expression, compared to the two other serotypes recommended to target brain cells (1 and 5). This is in accordance with the robust serotype 8 rAAV-mediated gene expression that has been observed and employed extensively non-optogenetic construct delivery to astrocytes in the rodent CNS. For example, Pignataro *et al* studied rAAV8 for astrocyte-specific delivery of therapeutic genes in the CNS [30], while Klein et al [31] demonstrated the use of AAV serotypes 8, 9, Rh10, and Rh43, with transduction efficiencies similar to that of AAV8. AAV8 and AAV7 vectors were also used for enhancing gene transfer efficiency in the murine striatum and in an orthotopic glioblastoma tumor model [32]. Interestingly, a synthetic serotype AAVDJ8 (generated by modifying specific residues of AAV8), displayed more tropism in astrocytes compared to other serotypes [33]. It is noteworthy that use of the GFAP promoter resulted in predominantly astrocytic transduction (~88%) with AAV8 [34] and there is high penetrance and almost complete specificity of gene delivery using AAV8-GFAP constructs to astrocytes [35]. Future work building on our study can characterize and quantify via immunohistochemistry, the percentage of each cell type transduced by serotypes of optogenetic construct employed in this study.

Recently, Schober *et al* [36] evaluated the transduction efficiency of serotypes 2 and 6 in non-optogenetic constructs in the rat brain cortex (*in vivo*) and found that serotype 6 efficiently transduced astrocytes. The implications of these modifications and the evaluation of a wider range of serotypes of the optogenetic construct targeting astrocytes is a future direction of this study. We recognize a gap in literature wherein, there is no systematic evaluation available for elucidating the most efficient AAV capsid serotype for the optogenetic construct targeting astrocytes in the rat brain cortex. Here we aimed to provide a foundation for the characterization of the optogenetic construct AAV-GFAP-ChR2(H134R)-mCherry, specifically for the light-induced intervention of astrocyte activity thereby establishing future robust control and mediation in several neurological disorders.

## Acknowledgements

This study was supported by the Wallace Coulter Foundation BME SEED grant - Florida International University (FIU) and Dissertation Year Fellowship (DYF) from the University Graduate School at FIU (Lakshmini Balachandar). The authors would like to sincerely thank Dr. Ranu Jung, and Dr. Timothy Allen for their help with equipment employed in this study and UNC Vector Core for providing the viruses used in this study. The authors would also like to thank Ms. Carolina Moncion for her feedback on the manuscript.

